# Human foot outperforms the hand in mechanical pain discrimination

**DOI:** 10.1101/2023.10.10.561422

**Authors:** Kevin K. W. Ng, Odai Lafee, Otmane Bouchatta, Adarsh D. Makdani, Andrew G. Marshall, Håkan Olausson, Sarah McIntyre, Saad S. Nagi

## Abstract

Tactile discrimination has been extensively studied, but mechanical pain discrimination remains poorly characterised. Here, we measured the capacity for mechanical pain discrimination using a two-alternative forced choice paradigm, with force-calibrated indentation stimuli (Semmes-Weinstein monofilaments) applied to the hand and foot dorsa of healthy human volunteers. In order to characterise the relationship between peripheral neural and perceptual processes, we recorded singleunit activity from myelinated (A) and unmyelinated (C) mechanosensitive nociceptors in the skin using microneurography. At the perceptual level, we found that the foot was better at discriminating noxious forces than the hand, which stands in contrast to that for innocuous force discrimination, where the hand performed better than the foot. This observation of superior mechanical pain discrimination on the foot compared to the hand could not be explained by the responsiveness of single primary afferents. We found no significant difference in the discrimination performance of either the myelinated or unmyelinated class of nociceptors between skin regions. This suggests the possibility that other factors such as skin biophysics, receptor density or central mechanisms may underlie these regional differences.

**Significance Statement:** Standard clinical practice for diagnosing neuropathies and pain disorders often involves assessing thresholds for pain or light touch. The ability to discriminate between different stimulus intensities is a separate but equally important sensory function, however this is not typically assessed in the clinic, and so studying this may provide insights into pain signalling mechanisms. Here, we investigated the ability of healthy individuals to discriminate between different forces of painful indentation. We found that the foot was better at this than the hand. This difference could not be explained by the firing activity of peripheral nociceptors (pain-signalling neurons) between the two regions, suggesting that mechanisms other than nociceptor sensitivity are involved.

## Introduction

Mechanical pain perception is considered a function of myelinated mechano-nociceptors, primarily the small-diameter, thinly myelinated (Aδ) mechano-nociceptors (Rolke et al., 2006), with recent research also indicating a contribution from the large-diameter, thickly myelinated (Aβ) mechanonociceptors (Nagi et al., 2019). Pain intensity ratings are widely used in both experimental and clinical settings. However, one caveat with pain ratings is that, while they may provide some insight into discriminative ability, measures for discrimination (or difference) thresholds, such as just noticeable difference (JND) or the Weber fraction (Holway and Pratt, 1936) cannot be easily determined from ratings. Establishing the Weber fraction provides a measure for sensory discrimination which can be compared across different conditions and modalities (Norwich, 1987). The processes of detection and discrimination serve distinct functions and may underlie different neural mechanisms, as suggested in studies on touch (Romo et al., 2008; Kim et al., 2014) and vision (Mazor et al., 2020; Schöpper et al., 2020). While detection thresholds are widely used, exploring pain discrimination may offer additional insights into the neural pathways involved in acute pain signalling.

In the current study, we used forced-choice psychophysical tests to investigate the human perceptual capacity to discriminate between innocuous and noxious indentation forces. The primary advantage of using a forced-choice approach is to overcome the bias, which would otherwise be introduced due to differences in response criteria between participants during scaling (Clark and Clark, 1980). We also compared the discrimination performance between hand and foot dorsa since the resolution of the somatosensory system is not constant across skin sites. For example, the spatial acuity for pain in the glabrous skin of the hand follows a proximal-to-distal gradient, with the fingertip being the area of highest acuity, whereas in the hairy skin of the upper limb, nociceptive two-point discrimination performance decreases in a proximal-distal direction (Mancini et al., 2014). Thus, it is of interest to compare discrimination performance between skin sites and body domains.

Using the forced-choice psychophysical method, we found that the capacity for discriminating noxious mechanical forces is significantly better in the foot than the hand. To explore whether this regional difference could be explained by different sensitivity of primary afferent nociceptors, we performed microneurography to record from myelinated (A) and unmyelinated (C) nociceptors innervating hand and foot dorsa. We found no difference between the hand and foot in the discrimination performance of either class of nociceptors, suggesting that a mechanism other than individual nociceptor sensitivity underlies the observed perceptual difference.

## Materials and Methods

We measured the psychophysical capacity to discriminate mechanical indentations of different forces applied to the dorsum of the hand and the foot in two psychophysical experiments. In the first experiment, high-intensity forces spanning a range of 100–3000 mN were used, targeting the noxious range of mechanical forces in which nociceptors display selective tuning and are rated as painful (Nagi et al., 2019). In the second experiment, low-intensity forces spanning a range of 6–80 mN were used, targeting the innocuous range of mechanical forces in which touch receptors display selective tuning (Middleton et al., 2022) and are clearly perceptible but not painful. To acquire neural data, we used the *in vivo* electrophysiological technique of microneurography (Vallbo, 2018) to record from single axons in the radial and peroneal nerves of awake participants. This was performed as a separate third experiment, using a subset of forces from both intensity series.

### Participants

For each of the two psychophysical experiments, we recruited 20 naïve healthy participants (noxious force range: 10 females, 21–33 years; innocuous force range: 6 females, 18–40 years). One participant took part in both experiments. For microneurography, we conducted new recordings with a separate group of 36 healthy participants (18–47 years). This study was approved by the Swedish Ethical Review Authority (2017/485-31 and 2020-04426), and the South West – Frenchay (20/SW/0138) and Liverpool John Moores University (14/NSP/039) research ethics committees. Informed consent was obtained from all participants in writing according to the revised Declaration of Helsinki.

### Equipment

A standardised set of Aesthesio nylon monofilaments (DanMic, San Jose, CA, USA), also termed von Frey hairs, was used to deliver innocuous and noxious mechanical stimuli. These filaments have different lengths and diameters based on the Semmes-Weinstein monofilament set, providing a linear scale of perceived intensity (Weinstein, 1993). The sizes of the monofilaments (1.65–6.65) correspond to a logarithmic function with equivalent forces ranging from 0.08 to 3000 mN (corresponding to pressures of 2.53 to 292 g/mm^2^). The monofilaments were applied manually (handheld), perpendicular to the test sites until they bent, with a contact time of approximately 1 s (Fig. 1A). The experimenter (O.L.) was trained to reliably apply the monofilaments in the intended way so that the filament always bent, and the tip did not slip along the skin. If hairs were visible on the test sites, they were removed by gently shaving the skin before applying the monofilaments (Cole et al., 2006). Participants were blind to visual cues by placing pillows to obstruct their field of view.

**Figure 1.**
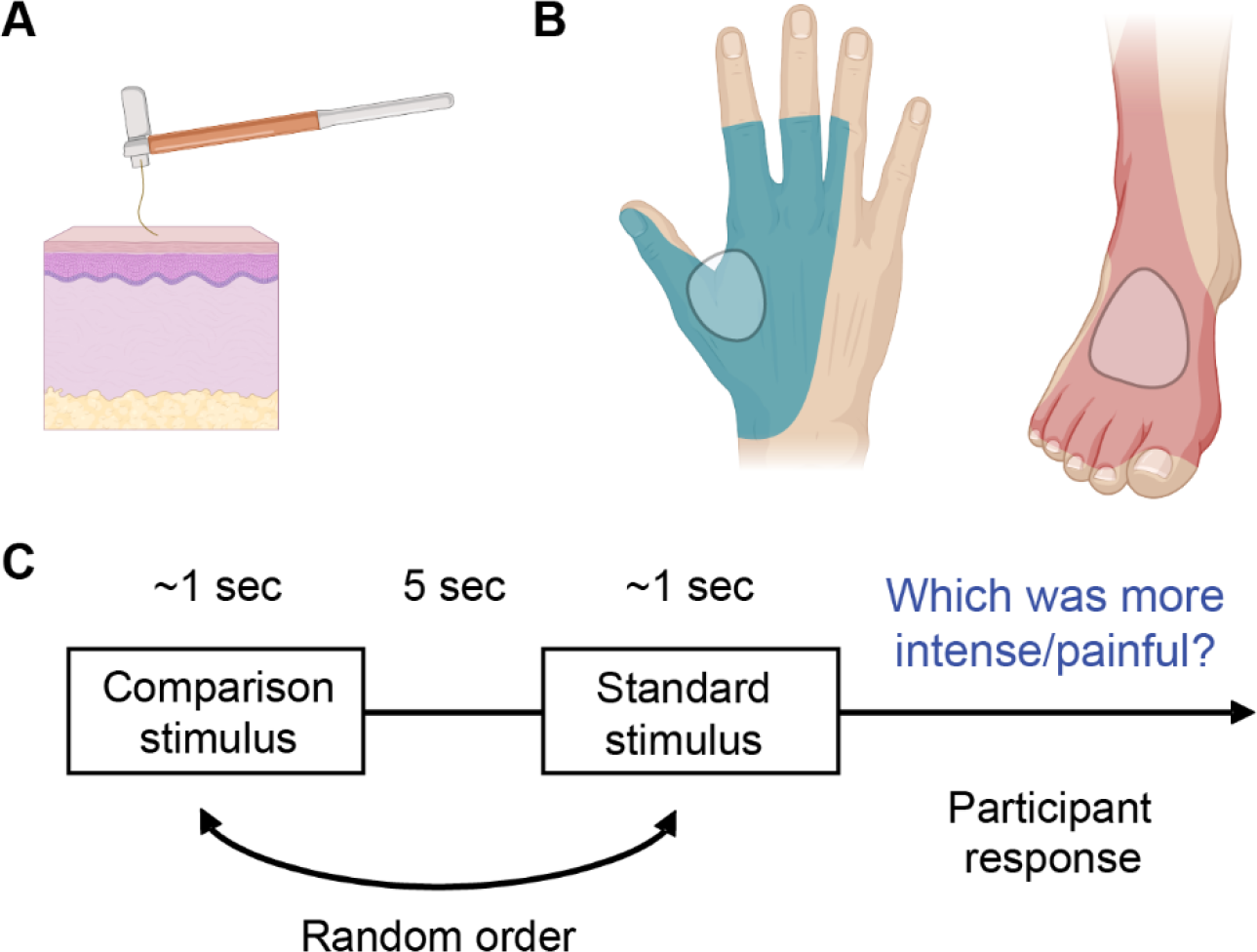
Schematic of the psychophysical experiment. (A) A von Frey filament is applied to the skin until it bends to deliver the target force. (B) The shaded skin regions indicate the innervation area of the radial and superficial peroneal nerves. The circled regions within this represent the stimulation sites where the monofilaments were applied. Images created with BioRender.com. (C) The two-alternative forced choice (2AFC) paradigm used for participants to judge which of the stimuli in the pair was perceived as “more intense” (with the innocuous forces) or “more painful” (with the noxious forces).

In microneurography experiments, LabChart software was used to process data acquired from a PowerLab 16/35 data acquisition system (ADInstruments, Sydney, Australia). An insulated high-impedance tungsten microelectrode (FHC, Bowdain, ME, USA) was inserted under real-time ultrasound (GE Healthcare, Chicago, IL, USA) guidance into the radial nerve proximal to the elbow or the superficial peroneal nerve proximal to the ankle. The reference (uninsulated) microelectrode was inserted just under the skin near the insertion point of the recording microelectrode. Neural activity was amplified using a headstage in conjunction with a low-noise high-gain Neuro Amp EX amplifier (ADInstruments).

### Force discrimination task

To determine the difference threshold on the hand and foot dorsa, i.e. the radial and peroneal territories respectively (Fig. 1B), we used a two-alternative forced choice (2AFC) psychophysical procedure in which two mechanical forces (a standard stimulus and a comparison stimulus) were presented successively in each trial, and the participants were asked to judge which stimulus was “more painful” (with the noxious force experiment) or “more intense” (with the innocuous force experiment) (Fig. 1C). The standard stimulus was always the same force within each experiment, and the comparison stimuli varied in force.

In the innocuous force series, the standard stimulus was 20 mN, and the comparison stimuli were 6, 10, 14, 20, 40, 60 and 80 mN (three stronger, three weaker and one equal to the standard stimulus). In the noxious force series, the standard stimulus was 600 mN, and the comparison stimuli were 100, 150, 260, 600, 1000, 1800 and 3000 mN. These intensities were chosen based on the psychophysical pain ratings in response to von Frey stimulation from previous work (Nagi et al., 2019). In a pseudorandom sequence, each of the comparison stimuli was paired 10 times with the standard stimulus to obtain a reliable estimate of the proportion of responses rated greater than the standard stimulus. The standard stimulus was presented first on half of the trials and second on the other half of the trials, in a random order, to minimise bias. All stimuli were applied for ∼1 s at inter-stimulus intervals of 5 s.

The stimuli (standard and comparison) were applied at different locations within the radial and peroneal territories. These locations were chosen randomly after each trial to avoid receptor fatigue or sensitisation. For each participant, the stimuli were delivered in two separate sessions during the same experimental sitting. In each session, the stimuli were delivered either on the hand dorsum or the foot dorsum. The order of sessions was counterbalanced across participants. The order sequence and timing of stimulus application was guided by a custom Python script (code available at https://github.com/SDAMcIntyre/Expt_MonofilamentDiscrimination). Participants were provided with a computer mouse to choose the more intense (innocuous series) or the more painful (noxious series) stimulus within each pair, and all the responses were registered automatically by the same program.

### Unit identification in microneurography

Isolated single afferents were searched for by brushing the skin using a soft or course brush, while making small adjustments to the position of the microelectrode. The A and C fibres were distinguished based on differences in spike morphology and response latency, with the C fibres displaying a characteristic delayed response to stimulation. In the case of nociceptors and touch receptors, distinction was made based on their response to soft brushing. All touch receptors are highly sensitive to soft brushing of their receptive field, whereas mechano-nociceptors do not respond to a soft brush but they may respond to a coarse brush and almost always respond to a pinch (Nagi et al., 2019).

In the current study, we recorded data from 31 new units and incorporated data from 9 units previously published in Nagi et al. (2019), resulting in a total of 40 units evenly split between myelinated and unmyelinated nociceptors and upper and lower limbs. All nociceptors were soft brush-insensitive with relatively high von Frey activation thresholds (≥4 mN), and conduction velocities, where measured, were >30 m/s for A fibres, suggesting Aβ range, and ∼1 m/s for C fibres, in line with previous observations (Vallbo et al., 1999; Nagi et al., 2019; Yu et al., 2023). The von Frey forces delivered during the microneurography experiment were 4, 10, 20, 60, 100, 260, 1000 and 3000 mN.

### Data analysis

For the psychophysical data, curve fitting was performed using R and the quickpsy package (Linares and López-Moliner, 2016). Psychometric functions were constructed for each site (hand and foot) in every participant by plotting the proportion of responses called greater than the standard stimulus against the intensities of comparison stimuli obtained from the 2AFC experiments. Curves were fitted to the data using a logistic function. The difference threshold or JND was taken as one-half the difference between the values of the comparison stimulus at the 75% and 25% points on the psychometric function (Sharma et al., 2022). The Weber fraction, which represents the ratio of the JND to the standard stimulus, was then calculated for each site in each participant. Paired t-tests were used to compare the differences between both skin sites. Statistical analyses were performed using Prism software (Graphpad, San Diego, CA, USA)

The microneurography data were processed using LabChart, with action potentials or spikes identified from background noise using threshold crossing and template matching. We considered neural activity within the first 500 ms window following the first evoked spike for analysis, which has been shown previously as a reliable metric for nociceptor tuning to indentation forces. This time window has also been shown to be sufficient to achieve the target indenting force across a wide range, as confirmed through testing with electronic von Frey monofilaments that provide force readouts (Nagi et al., 2019). Furthermore, the reaction time to punctate tactile stimulation is ∼300 ms (Lele et al., 1954), indicating rapid signalling and information processing.

Curve fitting and analysis of the processed neural data was then completed using Prism (Graphpad, San Diego, CA, USA). We fitted a semi-log line to the data comparing von Frey forces and the firing rate of recorded afferents, where indentation force on the x-axis was logarithmic. For units where data were collected from repeated stimulus applications, the trial which provided the highest firing rate at that particular force was chosen for the purposes of curve fitting. To compare if the neural responses differed between the hand and foot sites, we performed an extra sum-of-squares F test between two models. This computes which of the two models provides a better explanation for the data: one where a single slope could be fitted to the pair of datasets being compared, and another where each dataset has their own slope.

## Results

### Discrimination sensitivity

We constructed psychometric curves for each participant based on the results from the 2AFC task in the noxious force (Fig. 2) and innocuous force (Fig. 3) range to calculate Weber fractions. A steeper slope indicates greater discrimination ability, i.e. lower Weber fractions.

**Figure 2.**
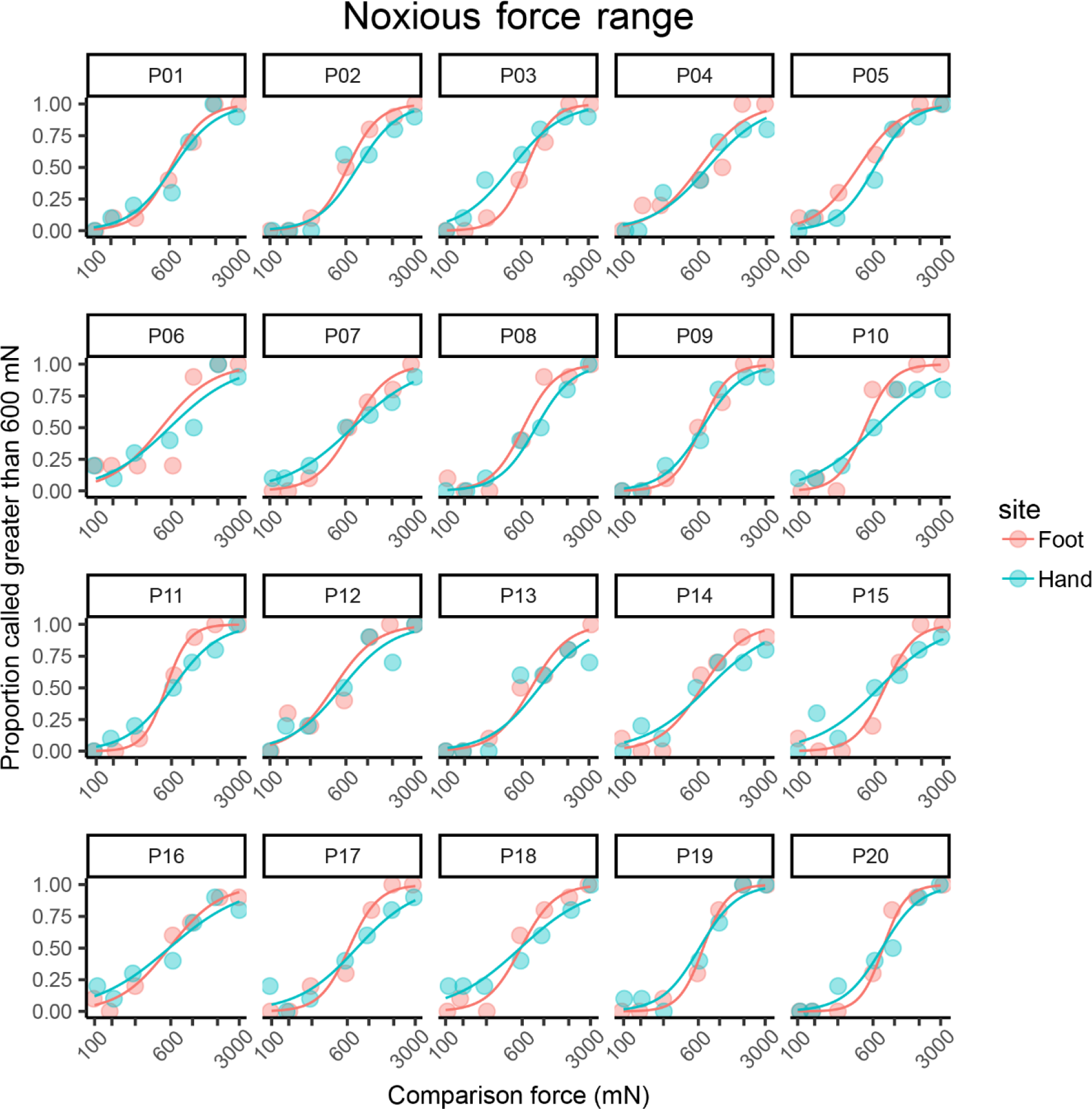
Psychometric function curves for mechanical force discrimination in the noxious force range (100–3000 mN) obtained from the psychophysical 2AFC task in the hand and foot of each participant (n = 20). Each data point was computed from 10 trials. Note the logarithmic scale on the x-axis.

**Figure 3.**
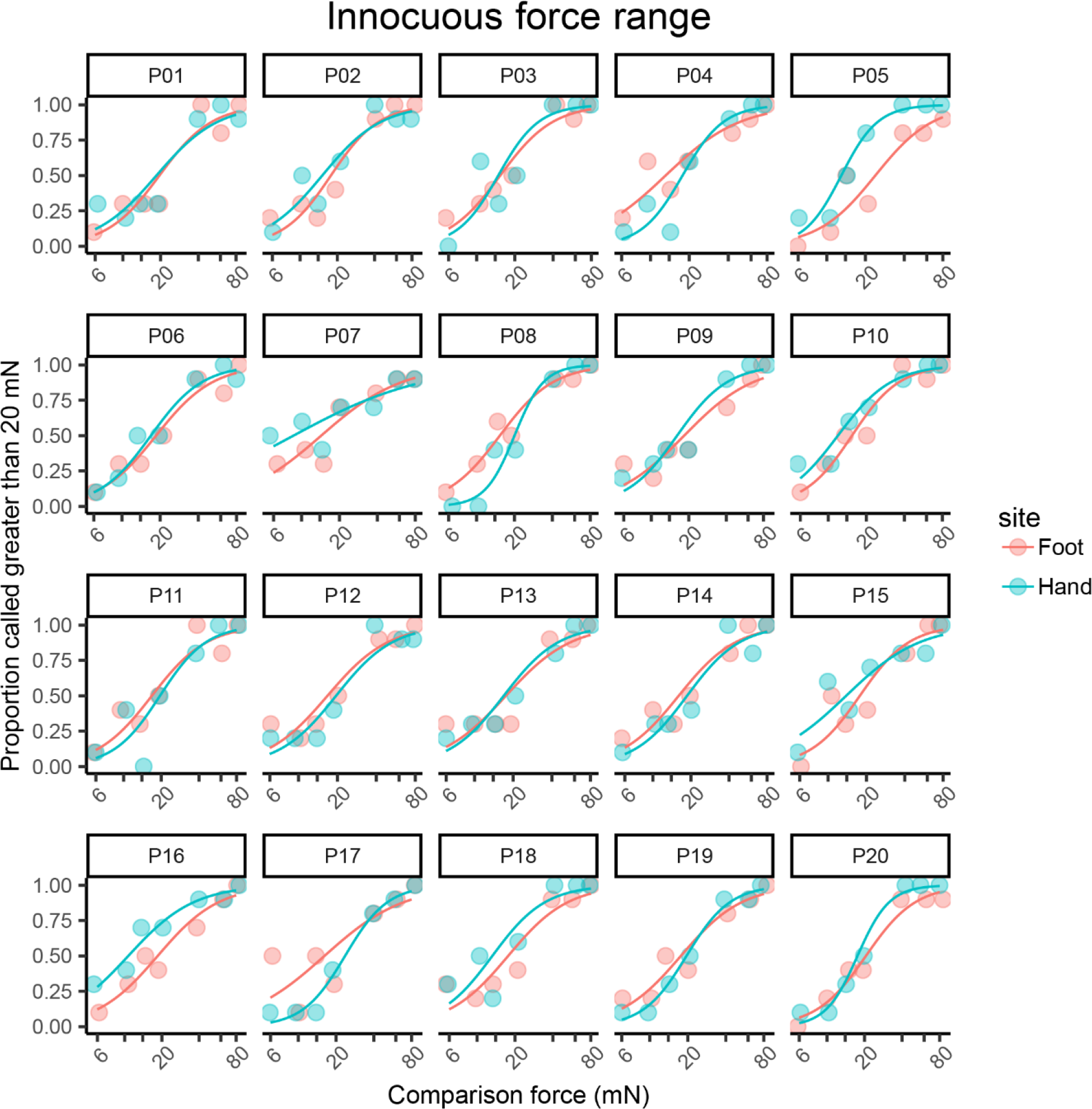
Psychometric function curves for mechanical force discrimination in the innocuous force range (6–80 mN) obtained from the psychophysical 2AFC task in the hand and foot of each participant (n = 20). Each data point was computed from 10 trials. Note the logarithmic scale on the x-axis.

The mean Weber fraction for discrimination of noxious mechanical stimuli was 0.88 (95% CI 0.78–0.99) in the hand and 0.52 (95% CI 0.46–0.58) in the foot. That is, at a force of 600 mN, a change of approximately 88% and 52% is required to be reliably perceived as more painful in the hand and foot, respectively. The Weber fraction in the foot was significantly lower (t(19) = 8.580, p < 0.0001) than that in the hand, as shown in Figure 4A. This contrasts with the results in the innocuous range (Fig. 4B), where discrimination performance in the hand (WF = 0.47, 95% CI 0.41–0.53) was better than that in the foot (WF = 0.57, 95% CI 0.53–0.62). This difference was also statistically significant (t(19) = 2.940, p = 0.0084).

**Figure 4.**
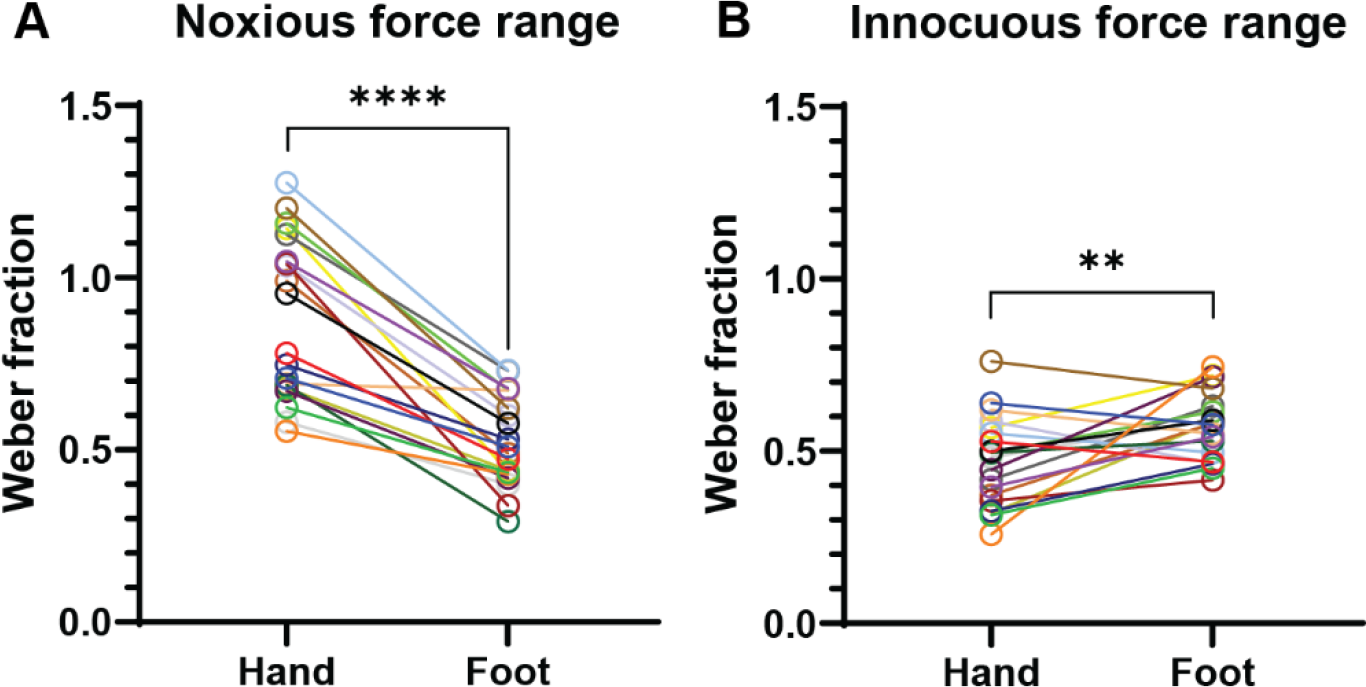
Within-participant comparison of Weber fraction between the hand and foot dorsa in both ranges of stimulation forces. (A) The foot is better at discriminating noxious mechanical forces than the hand. (B) The hand is better at discriminating innocuous mechanical forces than the foot. Each pair of circles connected by a line represents an individual participant (n = 20 each for innocuous and noxious force range experiments).

### Microneurography data

We recorded from nociceptors using microneurography and determined their mean discharge rates during the dynamic phase (500 ms onset) of indentation (Fig 5A). For the A nociceptors, fitting individual slopes for each skin site provides the following values: 13.33 (95% CI 9.50–17.16; R^2^ = 0.40) for the foot and 10.15 (95% CI 5.92–14.39; R^2^ = 0.25) for the hand (Fig 5B). However, the model where a single slope (11.82, 95% CI 9.00–14.65; R^2^ = 0.33) was fitted to both datasets can sufficiently explain the data, and so the slopes were not significantly different (F(1, 140) = 1.231, p = 0.2692). Similarly, when comparing the responses of the C nociceptors between the foot (slope = 8.23, 95% CI 6.51–9.94; R^2^ = 0.59) and hand (7.83, 95% CI 6.39–9.26; R^2^ = 0.66) (Fig 5C), no significant difference was found between the sites (F(1, 124) = 0.1244, p = 0.7249) as a single common slope (8.04, 95% CI 6.92–9.15; R^2^ = 0.62) could adequately fit both datasets.

**Figure 5.**
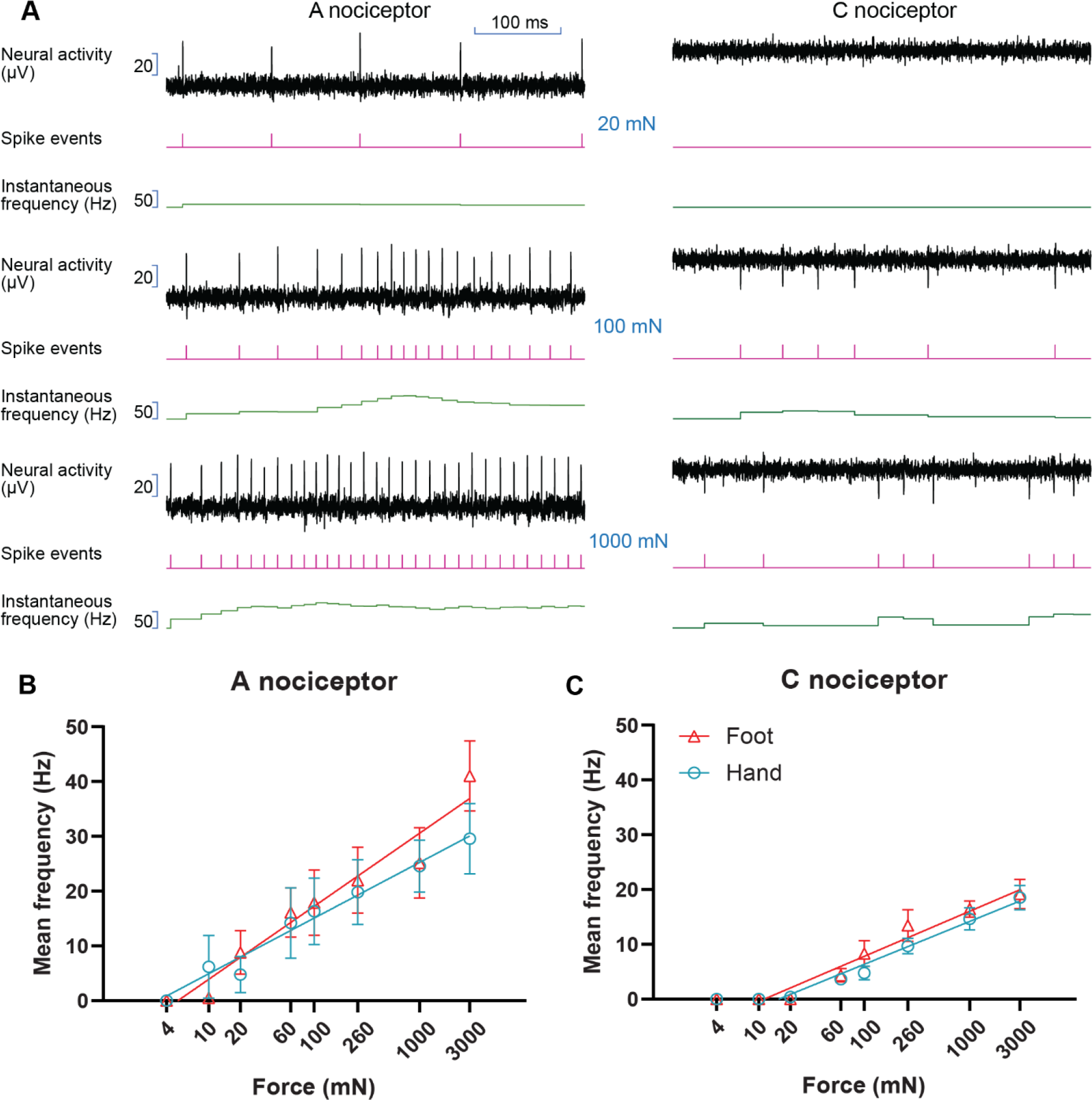
Responses of nociceptors during the dynamic phase (500 ms onset) of von Frey stimulation with different forces. (A) Recording traces of A and C nociceptors from the hand during the onset period at three different stimulation forces. (B) Comparison of mean firing rates of hand and foot A nociceptors (n = 10 units each site). (C) Comparison of mean firing rates of hand and foot C nociceptors (n = 10 units each site). Error bars represent SEM. Note the logarithmic scale on the x-axis.

## Discussion

In the current study, the human perceptual capacity to discriminate innocuous and noxious mechanical forces in the hand and foot dorsa was investigated. Our findings show that the foot is significantly better at discriminating noxious mechanical forces than the hand. In contrast, the hand is significantly better at discriminating innocuous mechanical forces. Our results align with the mechanical detection and pain thresholds (MDT and MPT, respectively) reported in the normative quantitative sensory testing (QST) dataset of the widely used German Research Network on Neuropathic Pain (DFNS), where the MDT is lower in the hand, and the MPT is lower in the foot (Rolke et al., 2006).

Both A and C mechanonociceptors display encoding of noxious indentation forces. The very fast conducting (Aβ-range) myelinated nociceptors have some unique features; for example, they exhibit much higher peak firing rates (up to 300 Hz) and much less propensity for fatigue during repeated stimulation compared to their unmyelinated counterparts (Nagi et al., 2019). In the current study, when we compared the responses between the two skin sites, we found no differences within the overall responses of either A or C nociceptors. Thus, the psychophysical differences between the skin sites cannot be explained by the response properties of individual nociceptors of either class, and other peripheral or central factors should be examined.

One possible explanation of the body region differences might relate to the innervation density. A better sensitivity for innocuous mechanical stimulation in the hand has been attributed to the higher innervation density of touch fibres in that region (Corniani and Saal, 2020). Quantification of intraepidermal nerve fibre density (IENFD), a method involving examination of skin biopsies for diagnosis of small-fibre neuropathies, has revealed that the hand dorsum has a higher density of small fibres than the foot dorsum (Ling et al., 2015). However, there is contrasting evidence with one study showing that the spatial acuity for heat pain was higher on the fingertips compared to the hand dorsum, despite the lower IENFD on the fingertips (Mancini et al., 2013). Nonetheless, both tactile A and C fibres have been demonstrated to have modulatory functions on pain signalling (Arcourt et al., 2017; Larsson and Nagi, 2022), which have a higher density in the upper limb (Löken et al., 2022), and this could potentially influence pain sensitivity (Liljencrantz et al., 2017).

Skin biomechanics is another important factor to consider, as it can vary between different anatomical sites. These variations could be due to differences in underlying anatomical structures (Biesecker et al., 2009) or skin thickness (Oltulu et al., 2018), which can influence the distribution of mechanical forces on the skin (Pawlaczyk et al., 2013). Because of this, the same mechanical force applied to different sites could potentially recruit a different number or class of nociceptors even if their innervation density would be the same between sites. In the current study, the psychophysical testing was conducted on skin sites without an underlying thick layer of fascia or muscle bulk. However, other factors contributing to biophysical differences cannot be ruled out.

Several central processing factors may also be relevant for the observed body region differences. Studies investigating the neural coding of non-painful indentation and vibrotactile stimuli suggest that the activity of several different classes of tactile afferents contribute to an overall percept of intensity, with each afferent class having different weights or contributions to the sensation (Cohen and Vierck, 1993; Muniak et al., 2007). It might be that pain perception also depends on the overall activity in the population, comprising inputs from different nociceptor classes or even tactile classes. However, selective electrical activation of individual Aβ nociceptors produces painful percepts at those same stimulus intensities at which non-painful percepts, such as pressure or vibration, are produced when Aβ tactile afferents are selectively activated (Nagi et al., 2019).

Another central factor influencing somatosensory performance is cortical magnification. The human primary somatosensory cortex (S1) contains fine-grained topographic maps that reflect nociceptive inputs from the skin (Mancini et al., 2012). It is possible that the degree of magnification of nociceptive signals in S1 may correspond to discrimination ability at certain skin sites, mirroring that observed with tactile acuity (Duncan and Boynton, 2007). Furthermore, several of the aforementioned factors may relate to the hand and the foot having evolved differently to have distinct functional roles (Hashimoto et al., 2013).

A limiting factor regarding our methodology is that the monofilaments we used had different diameters, meaning that the contact areas were different. Previous studies have reported that the probe size influences the perception of sharpness and MPTs (Greenspan and McGillis, 1991, 1994). For this reason, mechanical pain is usually tested using custom-made weighted pinprick stimuli that have a constant diameter (Rolke et al., 2006). However, we chose to use Semmes-Weinstein monofilaments to avoid damaging the receptive field of recorded afferents. Moreover, they are inexpensive, easy to administer, and widely used in clinical practice to screen for peripheral neuropathy (Berquin et al., 2010; Katon et al., 2013). Using these filaments, we managed to successfully measure both pain and touch discrimination with acceptable measurement variability and found significant differences.

That chronic pain remains a significant clinical challenge with limited treatment options may, in part at least, be due to our limited understanding of its underlying mechanisms (Schmelz, 2021). For instance, the primary focus of QST is on absolute thresholds (Krumova et al., 2012), and other psychophysical studies involving mechanical pain have also focused on measuring detection thresholds (Pfau et al., 2020; Suzuki et al., 2022). Thus, investigating discrimination sensitivity for pain and collecting normative data for this function may help expand our knowledge regarding the mechanisms that underlie acute pain signalling in humans.

## Author contributions

S.M., S.S.N., and O.L. developed the psychophysical protocol with input from H.O. O.L. performed the psychophysical experiments. O.L. and K.K.W.N. analysed the psychophysical data with input from S.M. O.B., S.S.N., A.G.M. and A.D.M. performed the microneurography experiments with assistance from K.K.W.N. K.K.W.N. analysed the microneurography data with input from S.S.N. and O.B. K.K.W.N. produced figures and wrote the manuscript with input from all other authors. All authors have read and approved the manuscript for submission.

## Data availability statement

Data associated with this study is available at https://osf.io/ne6pj/.

## Acknowledgements

We thank Warren Moore for collating some experimental information and Francis McGlone for providing the lab space where a subset of the microneurography experiments were conducted.

